# Discovery of novel multi-functional peptides by using protein language models and graph-based deep learning

**DOI:** 10.1101/2023.04.14.536982

**Authors:** Jiawei Luo, Kejuan Zhao, Junjie Chen, Caihua Yang, Fuchuan Qu, Ke Yan, Yang Zhang, Bin Liu

## Abstract

Functional peptides are one kind of short protein fragments that have a wide range of beneficial functions for living organisms. The majority of previous research focused on mono-functional peptides, but a growing number of multi-functional peptides have been discovered. Although enormous experimental efforts endeavor to assay multi-functional peptides, only a small fraction of millions of known peptides have been explored. Effective and precise techniques for identifying multi-functional peptides can facilitate their discovery and mechanistic understanding. In this article, we presented a novel method, called iMFP-LG, for identifying multi-functional peptides based on protein language models (pLMs) and graph attention networks (GATs). Comparison results showed iMFP-LG significantly outperforms state-of-the-art methods on both multifunctional bioactive peptides and multi-functional therapeutic peptides datasets. The interpretability of iMFP-LG was also illustrated by visualizing attention patterns in pLMs and GATs. Regarding to the outstanding performance of iMFP-LG on the identification of multi-functional peptides, we employed iMFP-LG to screen novel candidate peptides with both ACP and AMP functions from millions of known peptides in the UniRef90. As a result, 8 candidate peptides were identified, and 1 candidate that exhibits significant antibacterial and anticancer effect was confirmed through molecular structure alignment and biological experiments. We anticipate iMFP-LG can assist in the discovery of multi-functional peptides and contribute to the advancement of peptide drug design.

**Availability and implementation:** The models and associated code are available at: https://github.com/chen-bioinfo/iMFP-LG.

**Supplementary information:** Supplementary data are available online.

## 1 Introduction

Functional peptides are short amino acid (AA) fragments, usually ≤50AA, playing an important role in the regulation of a variety of biological functions, such as regulating hormones, neurotransmitters, and growth factors [29]. Because of their excellent selectivity, effectiveness, relative safety, and good tolerability in the regulation of a variety of biological functions, functional peptides have attracted tremendous attention in medicine [10]. Up to now, peptides have been discovered with a wide range of functions [8, 37], e.g. anti-microbial peptides, anti-cancer peptides, etc.. Simultaneously, a growing number of peptides have been confirmed to have multiple functions. For instance, some antimicrobial peptides are lethal to cancer cells [14]. Effective and precise techniques for determining the multiple functions of peptides can encourage their discovery and mechanistic understanding. Unfortunately, biological experiments in the study of peptide functions are time-consuming and expensive in both labor and materials.

At present, computational approaches are getting tremendous success in the discovery of peptide functions. As the development of machine learning techniques, the methodologies of these approaches have gone through three stages: conventional machine learning-based methods, deep learning-based methods and protein language model-based methods. Conventional machine learning-based methods (e.g. AVPpred [35], PredAPP [48], AIPpred [24], THPep [31]) identified peptides by employing SVM and RF algorithms based on various feature engineering techniques including PSSM matrix [2], physicochemical properties [16], pseudo amino acid composition (PseAAC) [30] and so on. Despite their effectiveness, conventional machine learning-based methods are often hard to generalize to other peptide datasets. Taking the advance of automatically extracted features and high performance hardware, deep learning-based methods (e.g. DeepACP [46], Deep-AmPEP30 [43], xDeep-AcPEP [5], and ITP-Pred [3]) stand out by employing different deep learning architectures including Convolutional Neural Network (CNN), Long Short Term Memory (LSTM) and their combination for distinguishing functional peptides. However, due to small datasets of functional peptides, these supervised deep learning-based methods encounter the challenge of learning robust peptide representation. Recently, benefited from unsupervised training and large corpus, pre-trained language models have emerged as a powerful paradigm in the field of natural language processing (NLP) [7, 26, 47]. Protein language models (pLMs) were also proposed and applied into peptide identification, such as antibacterial peptides [49], anti-microbial [23], signal peptides [34], bitter peptides [4]. Above studies are mono-functional peptide prediction methods. However, a growing number of peptides have been discovered with multiple functions.

In contrast to the discovery of mono-functional peptides, multi-functional peptide identification is a multi-label classification task which assigns a set of relevant functional labels to a peptide simultaneously. Multi-label classification is more complex due to the difficulties in capturing hidden connections of labels and resolving data imbalances. Several works have made a difficult endeavor in discovery of multi-functional peptides. Tang et al. [33] and Li et al. [19] identified multi-functional peptides by using a multi-label deep learning method, which combines CNN and RNN to extract peptide features and assigns function labels separately. PrMFTP [44] improved the performance of multi-functional therapeutic peptide identification by employing multi-scale CNN, attention-based biLSTM. Since the number of multi-functional peptides is fewer than mono-functional peptides and is extremely imbalanced, these approaches can’t generate a comprehensive peptide representation and have high false positive rate prediction. Besides, they predicted all function labels separately without considering their correlation. Pre-trained pLMs and graph attention networks (GATs) offer solutions to these problems. The pLMs pre-trained on millions of protein sequences [28] can capture dependencies between amino acids and biological properties. GATs have an ability to capture the complex relationship by computing attention coefficients between various objects [40]. The advantages of both pLMs and GATs can provide helpful insights for discovery of multi-functional peptides.

In this study, we develop a novel method, iMFP-LG, for discovering multi-functional peptides based on pLMs and GATs. To the best of our knowledge, iMFP-LG is the first approach that considers the association between function labels to identify multifunctional peptides by converting the multi-label problem to the graph node classification. The computational results show iMFP-LG outperforms state-of-the-art methods on both multi-functional bioactive peptide (MFBP) and multi-functional therapeutic peptides (MFTP) datasets. iMFP-LG is also interpretable by visualizing the distribution of peptide representations and motif patterns obtained by the pLM, and function relationships captured by the GAT. We employed the iMFP-LG model to establish a robust peptide discovery pipeline. Through this pipeline, eight novel candidate multi-functional peptides were screened out with high confidence from the UniRef90 database, which is an extensive collection of millions of known peptides. The further biological experiments show one peptide has remarkable bioactivity in terms of antimicrobial and anticancer functions. These results demonstrate the capability of iMFP-LG for discovery of novel multi-functional peptides.

## 2 Results

### 2.1 Overview of iMFP-LG

iMFP-LG is constructed for identifying multi-functional peptides, specifically multi-functional bioactive peptide (MFBP) [33] and multi-functional therapeutic peptides (MFTP) [44]. Both of them are collected from the literature by searching specific keywords in Google Scholar (**Figure 1A**). The architecture of iMFP-LG consists of two modules: peptide representation module and a graph classification module (**Figure 1B**). In the peptide representation module, protein language model (pLM) is responsible for acquiring high-quality peptide representations through multi-head self attention mechanism (**Figure 1C**). In the graph classification module, graph attention network (GAT) (**Figure 1D**) is employed to capture the interrelationships among various functional features. The nodes in the peptide function graph are function labels and the edges are the correlation between functions. The node features are projected by pLMs according to peptide sequences, while edge weights are learned by GATs. All node features are further tuned to integrate complex relationships of peptide function labels according to the corresponding edge weights. At last, the multi-functions of peptides are determined by performing a binary prediction based on the tuned node features. Besides, adversarial training is used to improve model robustness and generalization ability (**Figure 1F**). To discover novel multi-functional peptides, we established a robust pipeline based on the trained iMFP-LG model (**Figure 1E**).

**Figure 1:**
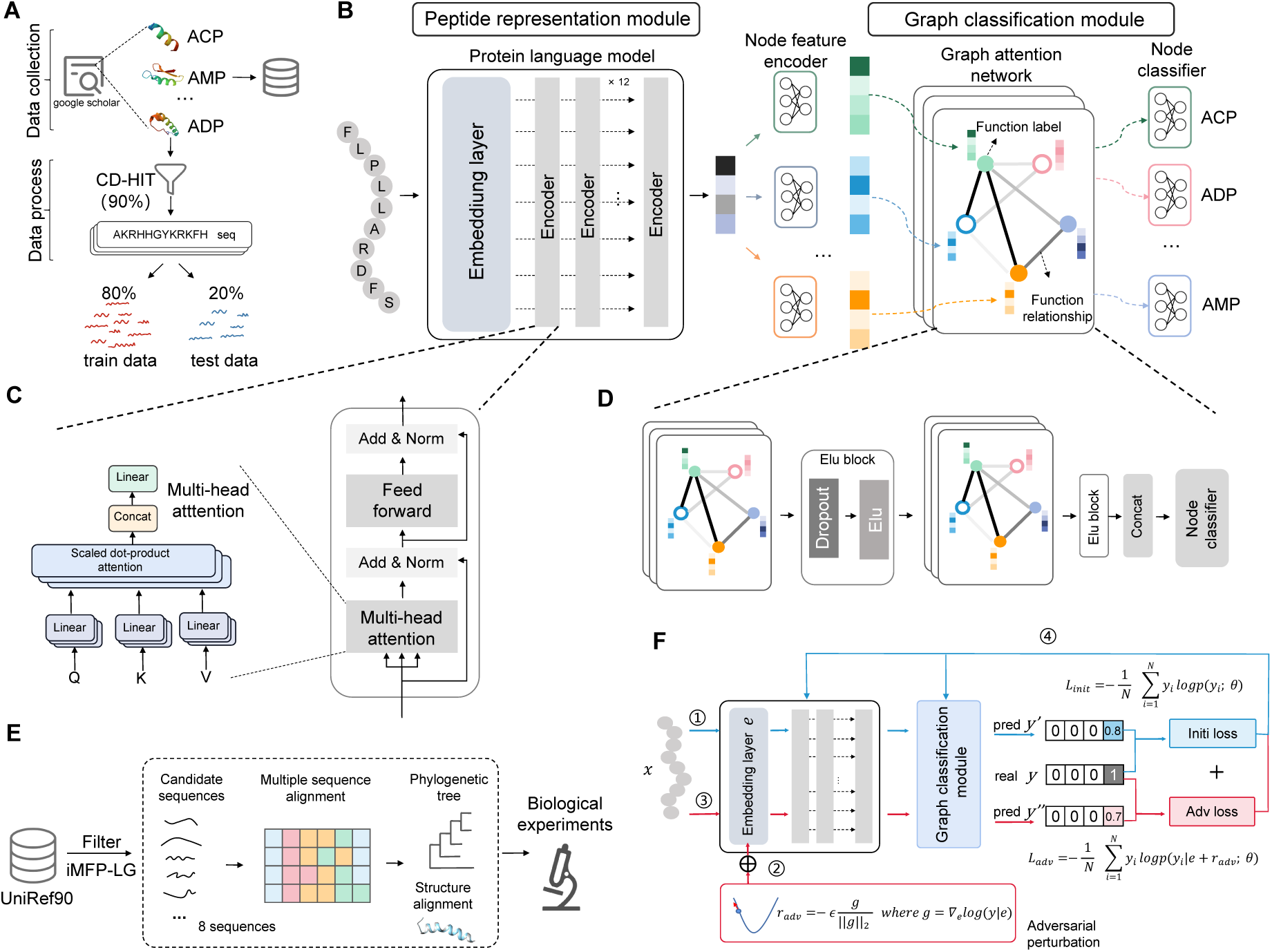
Illustration of the proposed iMFP-LG. (A) The collection and pre-process of multi-functional peptide datasets. (B) The framework of iMFP-LG, including a peptide representation module and a graph classification module. The peptide representation module employs a protein language model to extract high-quality peptide representation from its primary sequence. The graph classification module is composed of a node feature encoder, a graph attention network and a node classifier to learn the correlation of function labels. (C) The multi-head attention mechanism of an encoder layer in the protein language model. (D) Two stacked graph layers in the graph attention network. (E) The pipeline of peptide discovery by iMFP-LG. (F) The procedure of adversarial training.

### 2.2 Graph node classification framework improves discovery of multi-functional peptides

The discovery of multi-functional peptides is a multi-label classification task. To capture the complex relationship of multi-functions, we proposed a graph node classification framework. In this section, we evaluated the effectiveness of graph node classification framework on several widely-used sequence feature extraction methods, including four sequence composition features (CF) extracted by iFeatureOmega [6] e.g. Amino acid composition (AAC), Pseudo-amino acid composition (PAAC) [30], Distance-Pairs (DP) [21], Composition-Transition-Distribution (CTDD) [13], three deep learning-based methods (CNN, RNN, CNN+BiLSTM), and a pLM-based method (TAPE [27]).

We first evaluated the performance of above feature extraction methods with or without GAT on MFBP (**Figure2A**) and MFTP (**Figure2B**) datasets in terms of precision, coverage, accuracy and absolute true. We observed that all methods without GAT are surrounded by the corresponding methods with GAT in radar maps, which means the methods with GAT outperform the methods without GAT in terms of precision, absolute true, accuracy and coverage. These results demonstrate the proposed graph node classification framework can enhance the identification of multi-function peptides.

We also compared the performance of all competing methods on the both MFBP (**Figure2C**) and MFTP (**Figure2D**) datasets. The pLM without GAT outperforms the other features with or without GAT. And the performance of pLM is further improved by GAT, achieving the best performance with a precision of 0.777, coverage of 0.785, accuracy of 0.776, absolute true value of 0.767, and absolute false value of 0.082 on MFBP dataset and a precision of 0.721, coverage of 0.722, accuracy of 0.679, absolute true of 0.605 and absolute false of 0.032 on MFTP dataset. All specifically experimental results can be found in **Supplementary Table S1** and **S2**. These results indicate that pLMs can extract more comprehensive and high-quality features from peptide sequences compared to traditional features and other deep learning-based methods.

Thus, we constructed iMFP-LG by incorporating pLM as a feature extraction module and GAT as an identification module. To improve the generalization capability and avoid the over-fitting phenomenon, we also employed adversarial training to get better results in the final framework.

### 2.3 iMFP-LG outperforms the state-of-the-art methods

We compared our proposed method iMFP-LG with several state-of-the-art methods, including four conventional machine learning-based methods (CLR [12], RAKEL [36], RBRL [42] and MLDF [45]) and three deep learning-based methods (MPMABP [19], MLBP [33] and PrMFTP [44]). MPMABP and MLBP employed CNNs and RNNs for identifying multi-functional peptides. In addition to CNN and RNN, PrMFTP employed a multi-head self attention module to further optimize the extracted features for predicting multi-functional therapeutic peptides.

According to the performance comparison on the MFBP (**Figure 2E**) and MFTP (**Figure 2F**) datasets, iMFP-LG outperforms the state-of-the-art methods on both datasets in terms of all metrics, expect a comparable performance in terms of absolute false in MFTP dataset. When compared on the MFBP dataset, iMFG-LG achieves a precision of 0.797, coverage of 0.803, accuracy of 0.796, absolute true of 0.796, and absolute false of 0.078, greatly outperforming the state-of-the-art method MPMABP by 6.9%, 5.4%, 6.9%, 8.4% and 2.3% in terms of Precision, Coverage, Accuracy, Absolute true and Absolute false. When compared on the MFTP dataset, iMFP-LG achieves a precision of 0.730, coverage of 0.730, accuracy of 0.689, absolute true of 0.616, outperforming the state-of-the-art model PrMFTP by 3.1%, 6.1%, 3.8%, and 2.3% in terms of Precision, Coverage, Accuracy, Absolute true respectively. They achieves comparable performance in terms of Absolute false. The specifically computational results can be found in **Supplementary Table S3** and **S4**. Overall, iMFP-LG comprehensively outperforms all multi-functional peptide prediction methods.

**Figure 2:**
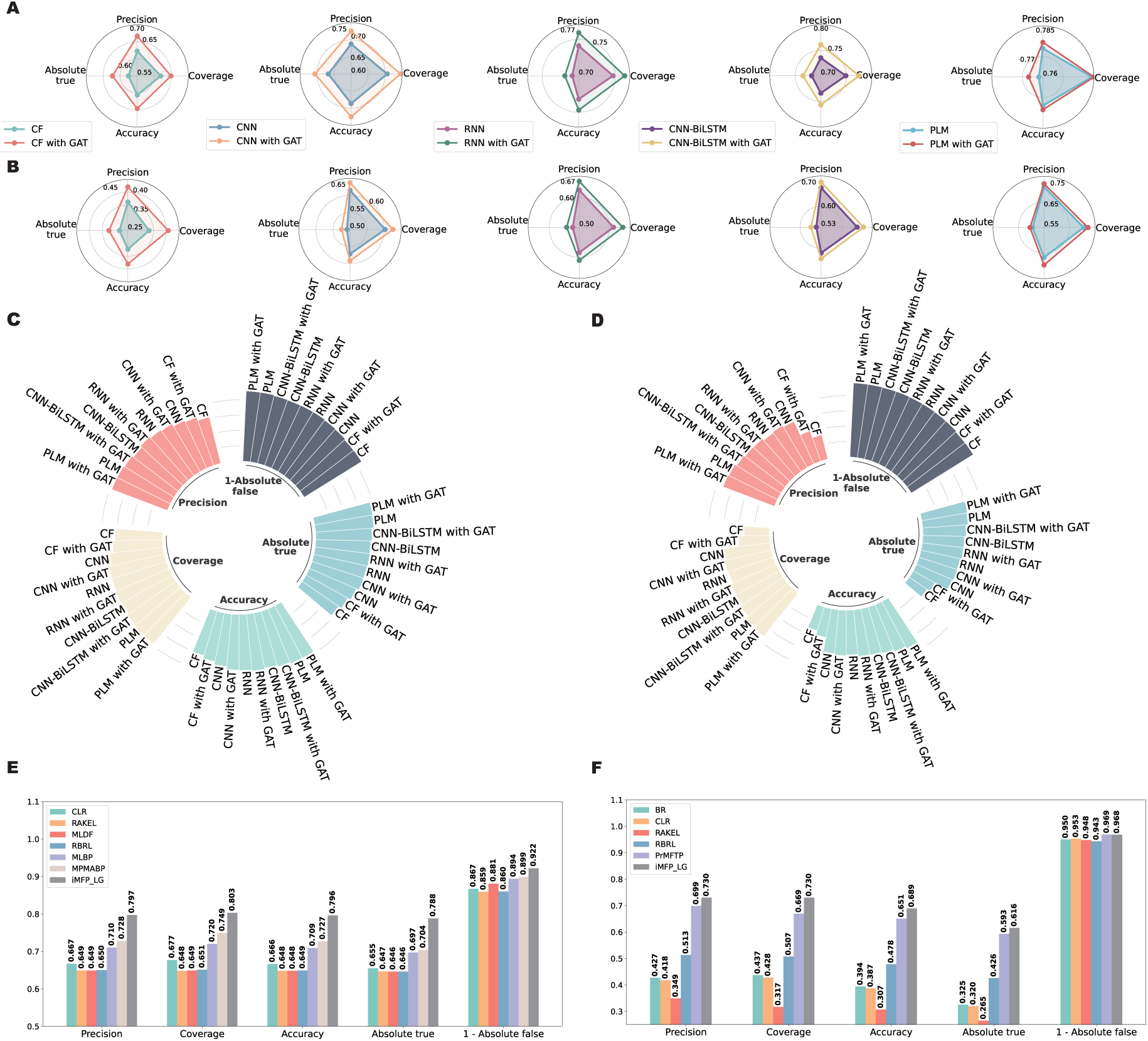
The performance comparison of different methods for identification of multifunctional peptides. (A) and (B) show the effect of GAT on different feature extraction methods in MFBP and MFTP datasets, respectively. (C) and (D) show the performance comparison of different feature extraction methods on MFBP and MFTP datasets, respectively. (E) and (F) are the performance comparison of our proposed method iMFP-LG and the state-of-the-art methods on the MFBP and MFTP datasets, respectively.

### 2.4 iMFP-LG is more sensitive to peptide categories with small size and multi-functions

As there are 5 types of bioactive peptide functions in MFBP and 21 types of therapeutic peptide functions in MFTP, we further investigated the performance of competing methods in each peptide category.

We compared the sensitivity (SEN) and specificity (SPE) of iMFP-LG with other state-of-the-art methods on all kinds of peptide functions. The SEN and SPE were calculated by considering peptides with a function as positive samples and other peptides without that function as negative samples. The comparison results on MFBP (**Figure 3A**) and MFTP (**Figure 3C**) datasets show that all competing methods have high SPE on both datasets, indicating that these models have a high negative sample recall in single function classification. But their SEN is low and differs largely. For the MFBP dataset in terms of SEN, iMFP-LG achieves the best results in prediction of ACP, ADP, AIP, AMP. Specifically, it has much higher SEN than the other two competing methods in two small categories ACP and ADP. For the MFTP dataset in terms of SEN, iMFP-

**Figure 3:**
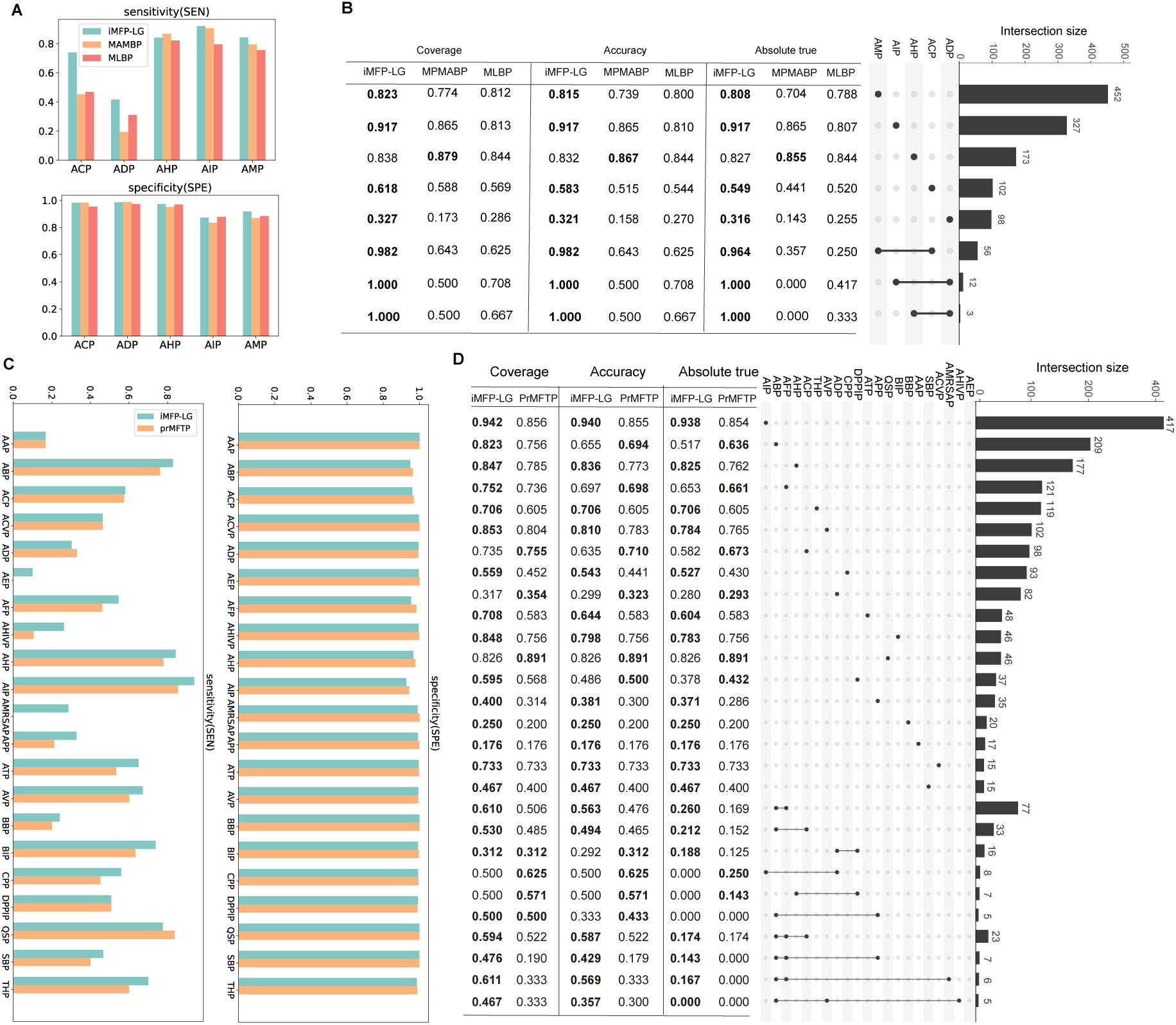
The performance comparison in each peptide category. (A) and (C) are sensitivity (SEN) and specificity (SPE) of different competing methods on the MFBP and MFTP datasets. (B) and (D) are the performance in mono-functional and multi-functional peptides on MFBP and MFTP, respectively. The upset plot in the right part shows the size of peptide categories in test datasets. For MFTP, we only show the peptide categories with size 5 and functions 3. The left part shows the performance in terms of coverage, accuracy and absolute true for corresponding peptide categories in the upset plot.

LG also achieves better results than PrMFTP in almost all peptide functions, except ADP and QSP. Interestingly, PrMFTP fails to distinguish the peptides in three small categories AEP, AHIVP and ADAMRSAP, nonetheless iMFP-LG greatly improves the prediction performance. These results demonstrate iMFP-LG is more sensitive to peptide categories with small size by taking the advantage of pLM to learn high-quality peptide representations.

We then investigated the performance with regard to mono-functional and multi-functional peptides on MFBP (**Figure 3B**) and MFTP (**Figure 3D**). For the MFBP dataset, iMFP-LG significantly outperforms the competing methods in almost all peptide categories, where for multi-functional peptides iMFP-LG achieves an absolute true of 1.00 in both ADP AIP and ADP AIP functions, and an absolute true of 0.964 in ACP AMP functions. For the MFTP dataset, iMFP-LG also achieves better performance than the competing method in most of peptide categories, especially in the multi-functional peptides. These results demonstrate iMFP-LG is more sensitive to peptide categories with multi-functions by taking the advantage of GAT to capture the complex function relationships.

### 2.5 Model ablation study

We then explored the contribution of each module to our proposed method through ablation experiments on two datasets. There are three important modules in iMFP-LG, including adversarial training, GAT, and pre-trained pLM. We built several variants of iMFP-LG with/without these modules.

**Table 1** and **Table 2** show the performance of iMFP-LG and its variants on MFBP and MFTP datasets. All variants keep consistent with the experimental settings except for learning rate in the ablation study. Because pLMs with random initialization are difficult to train at small learning rates, we set the learning rate to 1e-5 in the model w/o pretrain and 5e-5 for all others. As seen, removing any module from the proposed model reduces the performance. On the MFBP dataset, the performance of model w/o pretrain is the most deteriorated with an Accuracy of 0.752 and Absolute true of 0.733. The model w/o ad shows a reduction in performance after removing the adversarial training with an Accuracy of 0.776 and Absolute true of 0.767. Similarly, the performance of model w/o GAT is dropped with an Accuracy of 0.784 and Absolute true of 0.776. On the MFTP dataset, the performance of model w/o pretrain decreased drastically, and its Accuracy and Absolute true decrease by 7.1% and 6.9%, respectively. The performance of both model w/o GAT and the model w/o ad also decrease obviously in terms of all metrics.

**Table 1:**
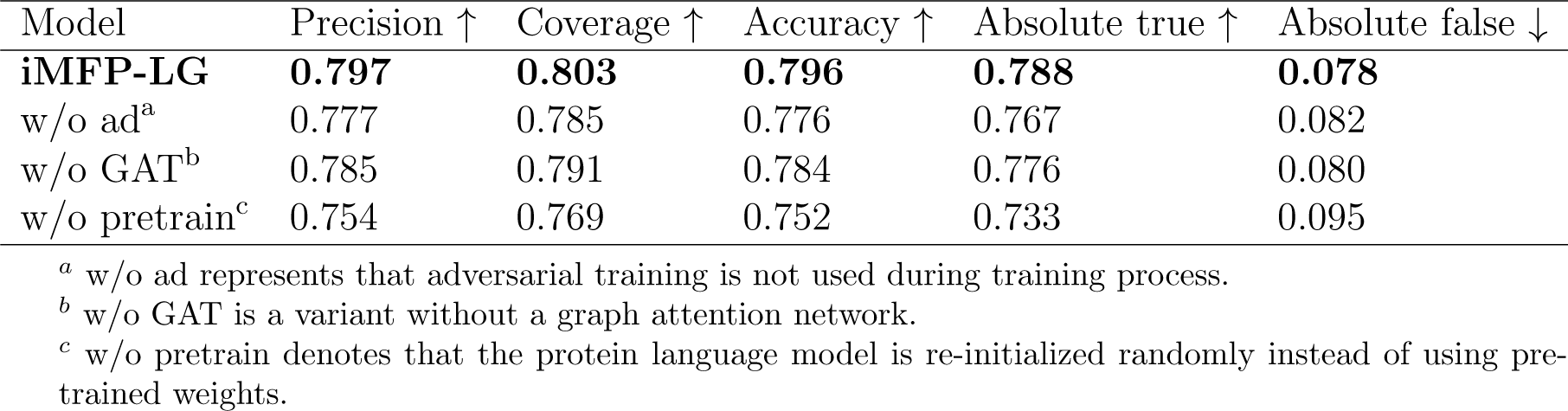
The performance of iMFP-LG and its variants on the MFBP dataset. The highest values are highlighted in bold.

| Model | Precision ↑ | Coverage ↑ | Accuracy ↑ | Absolute true ↑ | Absolute false ↓ |
| --- | --- | --- | --- | --- | --- |
| **iMFP-LG** | **0.797** | **0.803** | **0.796** | **0.788** | **0.078** |
| w/o ad<sup>a</sup> | 0.777 | 0.785 | 0.776 | 0.767 | 0.082 |
| w/o GAT<sup>b</sup> | 0.785 | 0.791 | 0.784 | 0.776 | 0.080 |
| w/o pretrain<sup>c</sup> | 0.754 | 0.769 | 0.752 | 0.733 | 0.095 |

<sup>a</sup> w/o ad represents that adversarial training is not used during training process.
<sup>b</sup> w/o GAT is a variant without a graph attention network.
<sup>c</sup> w/o pretrain denotes that the protein language model is re-initialized randomly instead of using pre-trained weights.

**Table 2:** The performance of iMFP-LG and its variants on the MFTP dataset. The highest values are highlighted in bold.

| Model | Precision ↑ | Coverage ↑ | Accuracy ↑ | Absolute true ↑ | Absolute false ↓ |
| --- | --- | --- | --- | --- | --- |
| **iMFP-LG** | **0.730** | **0.730** | **0.689** | **0.616** | **0.032** |
| w/o ad<sup>a</sup> | 0.721 | 0.722 | 0.679 | 0.605 | 0.032 |
| w/o GAT<sup>b</sup> | 0.709 | 0.705 | 0.667 | 0.598 | 0.033 |
| w/o pretrain<sup>c</sup> | 0.658 | 0.657 | 0.618 | 0.547 | 0.036 |

<sup>a</sup> w/o ad represents that adversarial training is not used during training process.
<sup>b</sup> w/o GAT is a variant without a graph attention network.
<sup>c</sup> w/o pretrain denotes that the protein language model is re-initialized randomly instead of using pre-trained weights.

Ablation studies reveal that all three modules have critical contributions to improve the performance of multi-functional peptides prediction, especially the GAT and pLM.

### 2.6 Interpretability of iMFP-LG

The intuition of iMFP-LG is to extract key sequence patterns via pLM and capture intricate relationships of multi-functions via GAT. Thus, we could unveil its decision process of how to assign multi-function labels to peptides by visualizing the distribution of peptide representations, sequence motifs and multi-function relationships.

All peptide representations were extracted by the pLM from the test dataset of MFBP and MFTP and visualized through tSNE [38]. We compared the distribution of peptide representations extracted by the pre-trained pLM on large protein corpus (**Figure 4A** and **4C**) and the fine-tuned pLM on multi-functional peptide datasets (**Figure 4B** and **4D**) on MFBP and MFTP, respectively. It is obviously to find that although the pre-trained model initially identifies different multi-functional peptides, the fine-tuned model exhibits more distinct ability to cluster peptides into categories with same functions. Interestingly, the peptide clusters with multiple functions lie in the middle of corresponding clusters of mono-functional peptides. Such as for the distribution in the MFBP (**Figure 4B**), the peptide cluster with ADP AIP function lies in the middle of the peptide cluster with ADP and peptide cluster with AIP. And the peptide cluster with ACP AMP function is also close to the cluster with ACP function and cluster with AMP function. The similar phenomenon is also observed in the distribution in MFTP (**Figure 4D**), such as the distribution of peptide clusters with (ABP, ACP, and ABP ACP) and (ADP, DPPIP, and ADP DPPIP). These results suggest that iMFP-LG has a capability to map peptides into a robust representation space, which is spatial interpretability with the continuity property: two similar multi-functional peptides should not be projected as two remote points in the representation space. This advance gives us an opportunity to explore the phylogenetic relationship of multi-functional peptides from mono-functional peptides.

**Figure 4:**
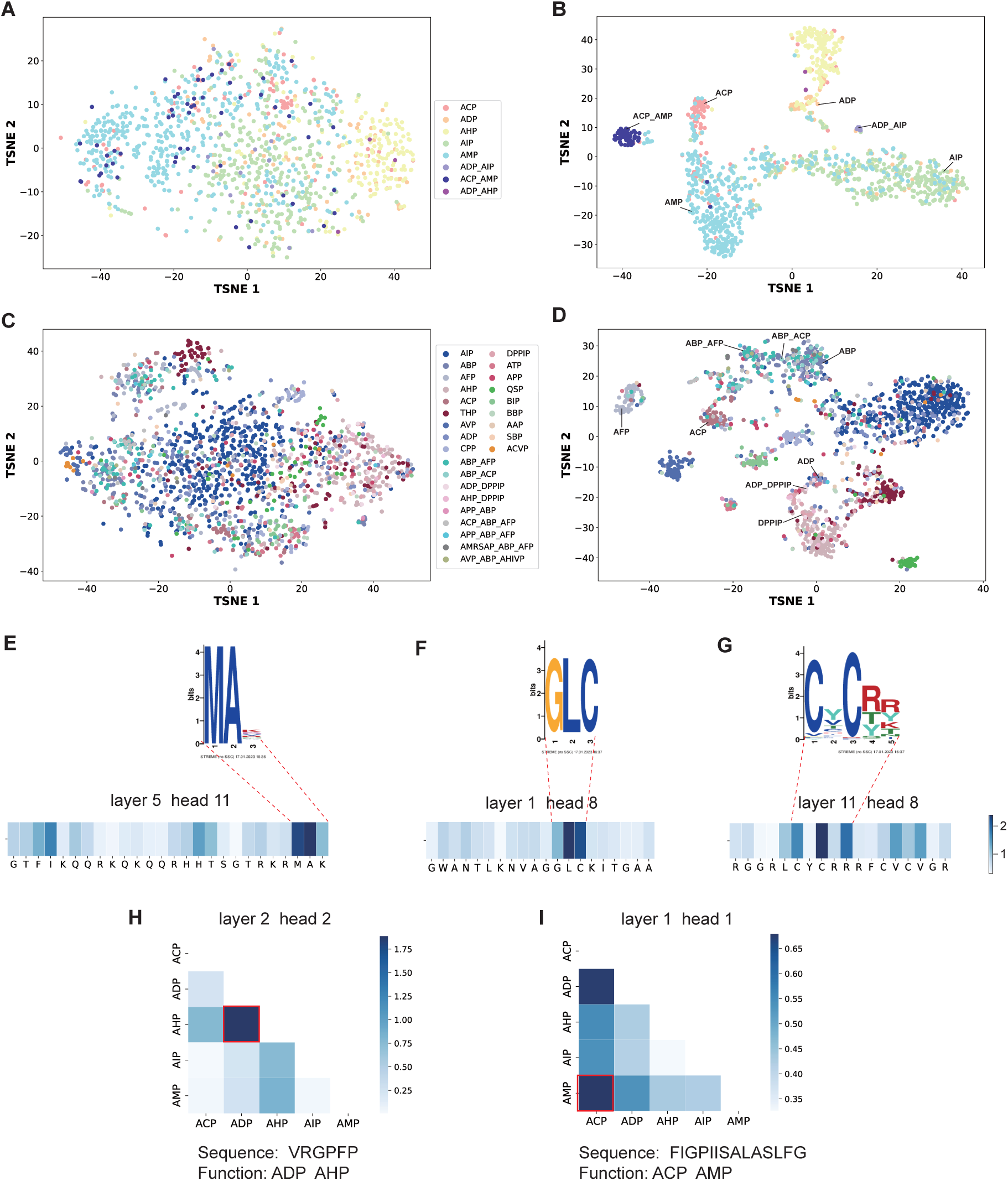
Illustration of the interpretability of iMFP-LG. tSNE visualization of distribution of peptide representations obtained by pre-trained and fine-tuned protein language models on MFBP dataset (A,B) and MFTP dataset (C,D), respectively. (E-G) describe 3 AMP cases whose sequence patterns captured by protein language model are matched with the motifs discovered by STREME. (H,I) show 2 multi-functional peptide cases that the learned graph node relationships are consistent with their true function labels.

To extract the sequence patterns from attention mechanism, we calculated the im-portance of each amino acid in a peptide sequence. Each sequence is transformed into an embedding by 12 attention heads in 12 layers, a total of 144 attention heads in the pLM. The weight *β_i,j_* in an attention head matrix indicates how much information from amino acid in position *j* needs to be used when computing the representation of amino acid in position *i*. We summed the attention of all amino acids to amino acid *j* as its effect 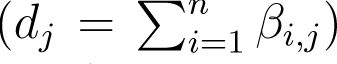 to the whole peptide sequence. Higher *d_j_* value suggests that amino acid *j* is more important in the peptide sequence. **Figure 4E-G** show 3 AMP cases that their patterns captured by attention mechanism are matched with the motifs discovered by STREME. The parameters of STREME is shown in **Supplementary Table S7**. We also used the attention visualization tool bertviz [41] to reveal the detail of the attention pattern (**Supplementary Figure S3**). These results suggest that iMFP-LG can find functional regions of peptides.

To interpret the intricate relationship captured by GAT, we visualized the graph attentions in 2 layers and 6 heads. The node relationship *r_i,j_* = *γ_i,j_* + *γ_j,i_* is calculated from graph attention matrix *γ*, in which each element *γ_i,j_* describes the weight of edge from node *i* to node *j*. We showed 2 multi-functional peptide cases to illustrate that the captured connections between graph nodes are consistent with true function labels. **Figure 4H** shows a node relationship captured by the layer 2 head 2. The edge between node ADP and node AHP has the largest attention score, which is the same as its true function label. **Figure 4I** shows another case that the edge between node ACP and node AMP has the largest attention score, which matches with its true function label. These results indicate that iMFP-LG can learn the relationship of different function labels.

### 2.7 Discovery of novel multi-functional peptides

Regarding to the outstanding performance of iMFP-LG on the identification of multi-functional peptides, especially on the MFBP dataset, iMFP-LG achieves an absolute true of 0.964 in ACP AMP functions, we employed iMFP-LG to screen novel candidate peptides with both ACP and AMP functions from millions of known peptides in the UniRef90.

Since UniRef90 contains 166,459,614 protein sequences, in order to focus on peptides, we filtered out any sequences *>* 40AA and *<* 4AA. This resulted in a dataset of 1,077,593 peptide sequences, which were then fed into iMFP-LG. To discover novel multi-functional peptides, we re-trained the iMFP-LG on the whole MFBP dataset with the same hyperparameters. To obtain candidate peptides with a high confidence, the classification threshold was set to 0.95 for both ACP and AMP functions. After removing duplicated peptides that appeared in MFBP dataset, we ultimately achieved 8 candidate peptides (**Figure 5A**). These peptide functions were further verified via searching their homologous sequences (**Figure 5B** and **Supplementary Figure S4**A) through multiple sequence alignment (MSA) [1] with candidate sequences as queries and peptides with ACP or AMP functions in MFBP dataset as targets. For these candidates that have ≥ 4 homologous sequences, we successfully constructed their phylogenetic trees, which demonstrate these candidates have evolutionary relationship to known ACPs and AMPs. We then predicted their structures by using ESMFold [20], and performed a structural alignment with their homologous sequences. The peptide UniRef90 P82904 exhibits exceptional alignment results(**Figure 5C**), but the peptides UniRef90 P83653 and UniRef90 B9W4V2 have no similar structures with their homologous sequences (**Supplementary Figure S4**B-C).

**Figure 5:**
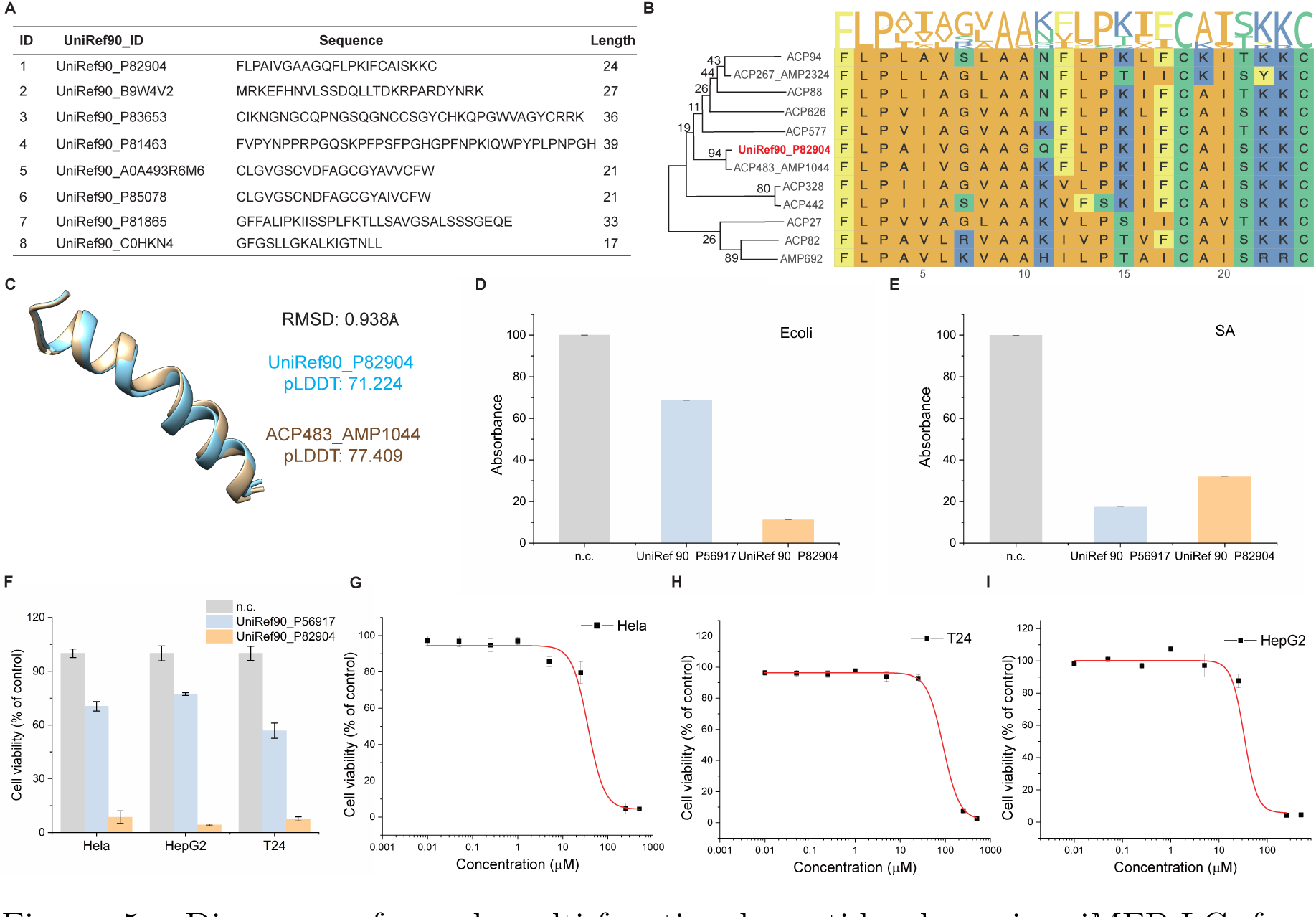
Discovery of novel multi-functional peptides by using iMFP-LG from UniRef90. (A) The candidate peptides with both anticancer and antimicrobial functions screened from UniRef90 by iMFP-LG. (B) Multiple sequence alignment and phylogenetic trees of candidate peptides UniRef90 P82904. (C) Structure alignment of the candidate peptide UniRef90 P82904 with its homologous sequence. (D-E) Bacterial inhibition effect of peptides UniRef90 P56917 and UniRef90 P82904 on E. coli and S. aureus bacteria strains. (F) Cytotoxic effect of peptides UniRef90 P56917 and UniRef90 P82904 on three T24, Hela, HepG2 cells. (G-I) Dose-dependent cytotoxic effect of peptides on Hela, T24, and HepG2 tumor cells.

In order to further assess the function of these three candidate peptides, we subsequently conducted biological experiments. A positive control (UniRef90 P56917) that has both AMP and ACP functions [17, 32] was random selected. Peptides at 500 *µ*M were added to cell culture for 24 hours, followed by MTT cytotoxicity assay. Three independent replicates were conducted. The results show that the peptide UniRef90 P82904 has excellent antibacterial effect on both E. coli bacteria and S. aureus bacteria (**Figure 5D, E**). The other two peptides UniRef90 P83653 and UniRef90 B9W4V2 have no antibacterial activity on both E. coli bacteria and S. aureus bacteria(**Supplementary Figure S4**D, E). To assess the anticancer activity of these peptides, the cell viability tests were performed on three kinds of human tumor cell lines, including bladder cancer (T24), cervical cancer (Hela) and liver cancer (HepG2). We found that the peptide UniRef90 P82904 has the strongest anticancer activity against all three tested cell lines (**Figure 5F**), and the peptide UniRef90 P83653 demonstrates strong anticancer activity against Hela cells, while having weak anticancer activity against T24 and HepG2 cells. The peptide UniRef90 B9W4V2 exclusively exhibits weak anticancer activity against T24 cells, and has no effect to other two tested cell lines (**Supplementary Figure S4**F). Subsequently, the peptide UniRef90 P82904 was chosen for a dose-dependence analysis in the 24 hours assay to evaluate its cytotoxic effect against three tumor cell lines by the standard MTT assay. As shown in **Figure 5G-I**, the cell viability decreased as low as 4.7%, 7.7%, and 4.3% at 250 *µ*gmL*^−^*^1^ against Hela, T24, and HepG2 cancer cells, thus demonstrating the promising capability for three cancer cell ablation. The outcomes of biological experiments are consistent with the computational screening of iMFP-LG, indicating that it has a strong potential to discover novel multi-functional peptides.

## 3 Conclusion

In this study, we proposed a novel method iMFB-LG for discovering multi-functional peptides. iMFB-LG converts multi-label prediction to graph node classification based on pLM and GAT. Comparison results in MFBP and MFTP showed that iMFB-LG significantly outperforms state-of-the-art methods, especially for small and multi-functional peptide categories. iMFB-LG is also interpretable through visualizing the patterns captured by attention mechanism of pLM and GAT. Subsequently, a peptide discovery pipeline was established based on iMFB-LG to screen novel multi-functional peptides. 8 candidate peptides with both AMP and ACP functions were discovered from the UniRef90 database. Further biological experiments of dose-dependence analysis demonstrate the promising anticancer and antibacterial activity of candidate peptides, indicating iMFB-LG has a strong potential to discover novel multi-functional peptides.

In future study, we plan to integrate function-related features, structure information and physic-chemical properties to strengthen the capability of graph nodes. Graph networks can be developed to delve further into the connections between various functional features. Although the success achieved in deep learning based prediction, experimental validating the predicted functions of peptide is still necessary for future study. In our opinion, other multi-label bioinformatics tasks can effectively use our technique as an extension.

## 4 Methods

### 4.1 Datasets

We evaluated our proposed method on two multi-functional peptide datasets, multi-functional bioactive peptide (MFBP) [33] and multi-functional therapeutic peptides (MFTP) [44]. Both of them were collected from the literature by searching specific keywords in Google Scholar. 80% of the data was randomly sampled as training data, and the remaining 20% was the test data (**Figure 1**A).

The MFBP dataset was collected by searching the keyword ‘bioactive peptides’ in the Google Scholar in June 2020. It contains 5986 bioactive peptides with 5 different functional attributes, including anticancer peptide (ACP), anti-diabetic peptide (ADP), anti-hypertensive peptide (AHP), anti-inflammatory peptide (AIP) and anti-microbial peptide (AMP). For each functional peptide category, CD-HIT [11] was applied to remove sequences with similarity greater than 90% in order to avoid redundancy and homology bias. Majority of bioactive peptides only have one type of activity, a few peptides have two types of activity jointly, and no peptide has more than two activities at the same time. The distribution of peptide categories is shown in **Supplementary Figure S1**.

The MFTP dataset was also collected by searching the keyword ‘therapeutic peptides’ in Google Scholar in July 2021. The collected data were pre-processed with following criteria: (1)Discard sequences containing non-standard amino acids; (2)Peptides longer than 50AA or shorter than 5AA are removed; (3)Delete peptides with fewer than 40 samples. After data processing, there are 9874 therapeutic peptides with 21 different functional attributes, including AAP, anti-bacterial peptide (ABP), anti-cancer peptide (ACP), anti-coronavirus peptide (ACVP), anti-diabetic peptide (ADP), anti-endotoxin peptide (AEP), anti-fungal peptide (AFP), anti-HIV peptide (AHIVP), anti-hypertensive peptide (AHP), anti-inflammatory peptide (AIP), anti-MRSA peptide (AMRSAP), anti-parasitic peptide (APP), anti-tubercular peptide (ATP), anti-viral peptide (AVP), blood-brain barrier peptide (BBP), biofilm-inhibitory peptide (BIP), cell-penetrating peptide (CPP), dipeptidyl peptidase IV peptide (DPPIP), quorum-sensing peptide (QSP), surface binding peptide (SBP) and tumor homing peptide(THP). The distribution of peptides in each type is shown in **Supplementary Figure S2**.

### 4.2 The proposed method iMFP-LG

iMFP-LG consists of two modules: peptide representation module and a graph classification module. The peptide sequences are first fed into the peptide representation module to extract high-quality representations, which are then transformed as node features by node feature encoders. The graph classification module is performed to fine-tune node features by learning the relationship of nodes. Finally, the updated node features are utilized to determine whether the peptides have corresponding function or not through node classifiers. Besides, the adversarial training also plays an important role to improve model robustness and generalization ability. The architecture of iMFP-LG and the adversarial training are introduced as follows.

#### 4.2.1 Peptide representation module

Here, a pre-trained pLM TAPE [27] is employed to generate the peptide representations. TAPE is a bidirectional encoder representation from transformer-based language model [7] constructed with a 12-layer Transformer encoder and 12 attention heads in each layer. The core of attention mechanism (**Figure 1C**) can be formulated as follows:

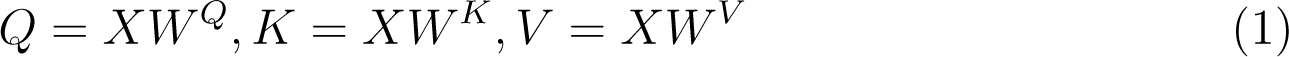

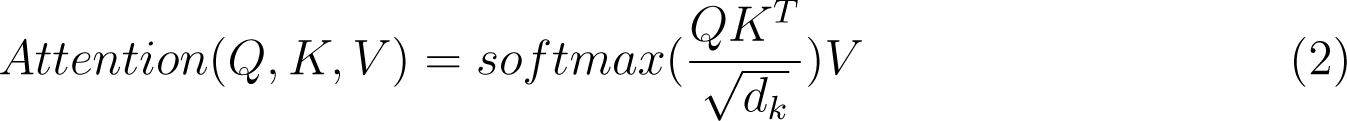

where *X ∈ R^L×d_m_^* is an embedding of a peptide sequence. The sequence embedding *X* will be transformed to Query matrix *Q ∈ R^L×d_k_^*, Key matrix *K ∈ R^L×d_k_^*, Value matrix *V ∈ R^L×d_k_^* by linear transformation and the *W^Q^, W^K^, W^V^ ∈ R^d_m_×d_k_^* are the learnable parameters of the attention layer. The attention scores are the normalized dot product of the key and query vectors in Eq. (2). The multi-head attention is an extension of the core of attention mechanism to catch the rich information from multiple projections, which can be formulated as follows:

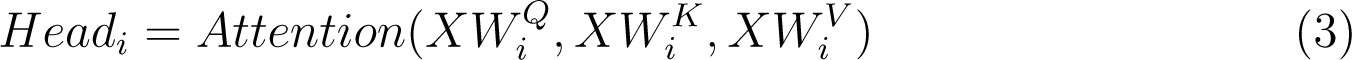

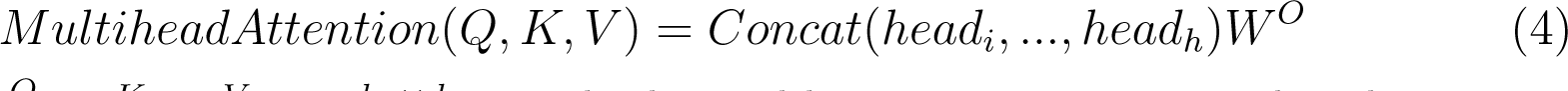

where the *W^Q^_i_, W^K^_i_, W^V^_i_ ∈ R^d_m_×d_k_^* are the learnable parameters in *Head_i_*. The output of the multi-head attention layer is obtained by concatenating the output of all attention heads, and then performing linear transformation with *W^O^ ∈ R^hd_v_ ×d_m_^*.

TAPE was pre-trained by masked-token prediction on the Pfam [9] corpus, which contains more than 31 million protein domains. The aim of masked-token prediction is to predict the randomly masked amino acids according to other amino acids in the protein. Thus, TAPE can model the general relationship of amino acids. In this study, we used the output of pooler layer in TAPE as peptide representations that are dimension 768.

#### 4.2.2 Graph classification module

Graph classification module is used to transform the multi-label classification problem into a graph node classification problem. It consists of three parts: node feature encoder, GAT and node classifier.

The node feature encoder constructs a linear transform layer for each graph node to convert the peptide representation as corresponding node representation, which can be formulated as follows:

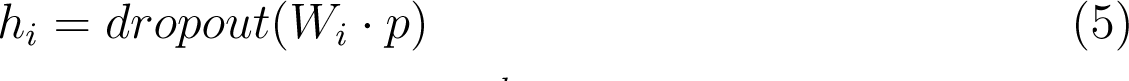

where *h_i_ ∈ R^d_p_LM^* represents the feature of node *i*, *p ∈ R^d_p_LM^* is the peptide representation, and *W_i_ R^d_p_LM ×d_p_LM^* is trainable parameters of the linear transformation layer. Specifically, a peptide representation with dimension of 768 was converted into 5 node representations for MFBP dataset and 21 node representations for MFTP dataset. All node representations are dimension 128.

The GAT is multi-head attention based graph neural networks for fine-tuning node representation by learning relationships among peptide functions. The graph nodes represent the different peptide functions and the edges represent the association between two functions. The node representation is updated by the GAT as follows:

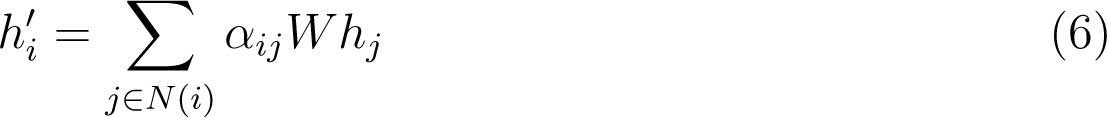

where *h^′^_i_* represents the updated node representation. *W ∈ R^d×d_p_LM^* is the transformation matrix. *α* depicts the attention matrix and *α_ij_* indicates the importance of the node *j* to node *i*. It can be calculated by:

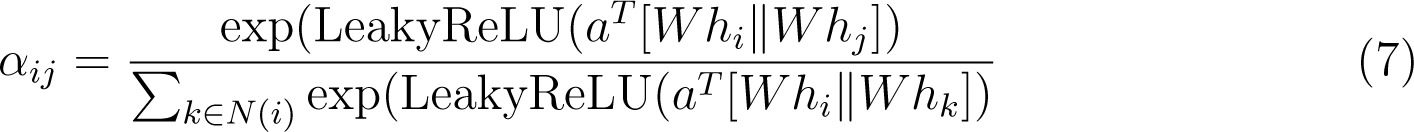

where *a^T^ ∈ R*^2^*^d^* is the parameter matrix of the attention mechanism, is the concatenation operation. Multi-head attention [39] is used in GAT. The final node features are concatenated by *K* independent attention heads.

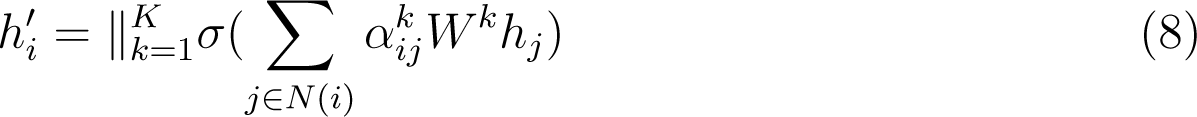

where *K* is the number of attention heads, *σ* is the activation functions, *∥* denotes concatenation operation, *α^k^_ij_* and *W^k^* are attention coefficients and weight matrix in *k*th attention mechanism. In this work, we created 2-layer fully-connected graphs among function labels, where all edge weights are initialized as 1 and each layer has 6 attention heads.

Node classifier is a set of binary predictors used for the final prediction of function labels. The updated node representations by GAT are fed into the corresponding binary predictors to predict whether the peptide has corresponding function or not. In this study, each binary predictor has a hidden layer and an output layer with sigmoid activation function, which can be formulated as follows:

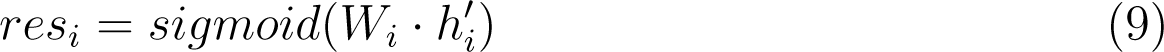

where *W_i_ ∈ R*^1^*^×d_p_LM^* is learnable in the hidden layer.

#### 4.2.3 Adversarial training

In order to improve the protein language representation’s capability and avoid the overfitting phenomenon, we employed an adversarial training strategy called Fast Gradient Method (FGM) [15, 25] during the training process.

FGM adds an adversarial perturbation to the embeddings of amino acids according to the updated gradients. The adversarial perturbation *r_adv_* can be defined as Eq. (10).

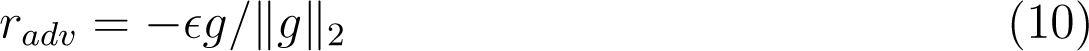

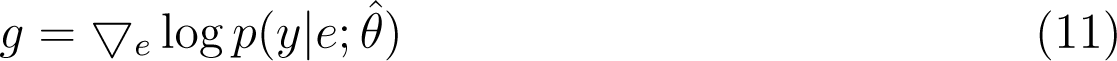

where *e* is the embedding of peptide sequences, *y* represents true function labels and *p*(*y e*; *θ*) represents conditional probability of *y* given *e*, *θ*^^^ is a constant set to the current parameters of the model and *ɛ* represents the shared norm constraint of adversarial loss. The goal of adversarial training is to minimize the original loss without perturbation and the adversarial loss with the adversarial perturbation (**Figure 1F**). The procedure of these two goals can be formulated as Eq. (12) and (13) respectively.

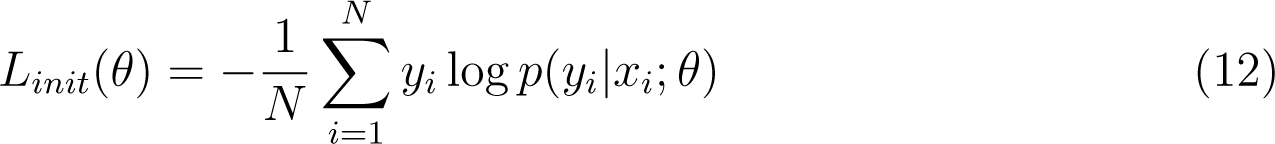

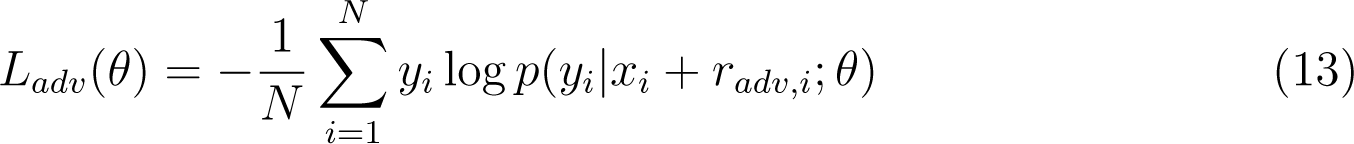

where *N* is the number of batch size, *x_i_, y_i_, r_adv,i_* are the input peptide sequence, label and perturbation on the *i*th sample in a batch and *θ* denotes the model’s parameters.

### 4.3 Evaluation metrics

In order to make a full and fair comparison with the state-of-the-art methods, we adopt 5 evaluation metrics, including precision, coverage, accuracy, absolute true, and absolute false. These metrics are defined as follows:

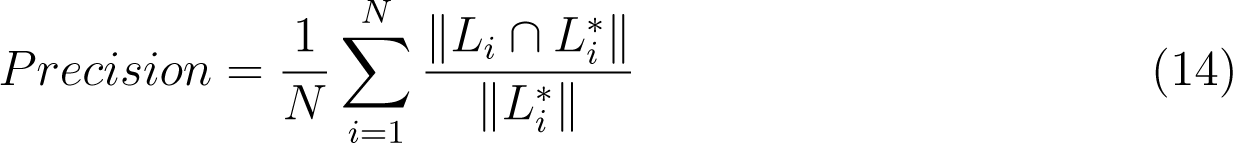

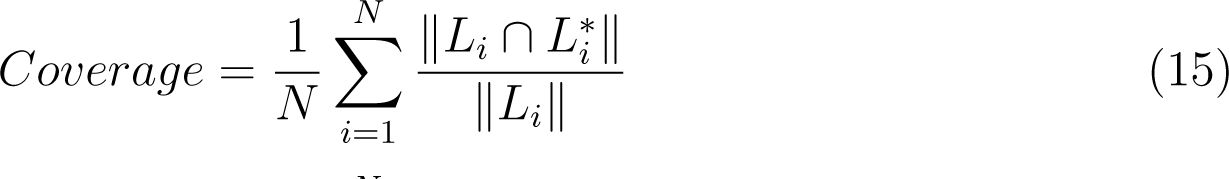

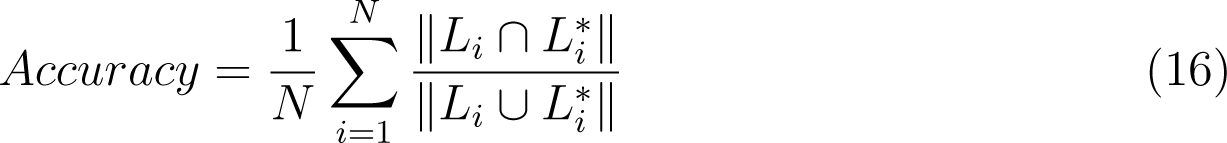

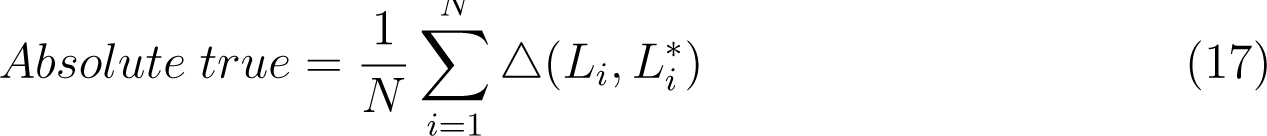

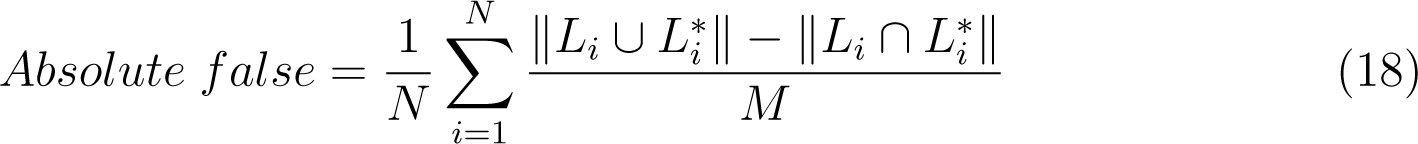

where *N* represents the total number of peptide sequences in the datasets, *M* denotes the number of labels, *∩* and *∪* is the intersect and union operation in the set theory, *S* represents the size of a set S, *L_i_* denotes the true label subset of the *i*th peptide sample, *L^∗^_i_* represents the predict label subset of *i*th sample by the classifier, and

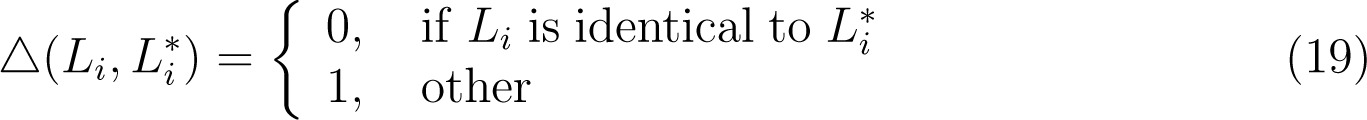

We also employed the sensitivity (SEN) and specificity (SPE) metrics [18, 33] to further compare the performance on each peptide function.

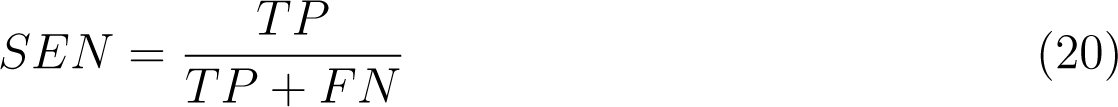

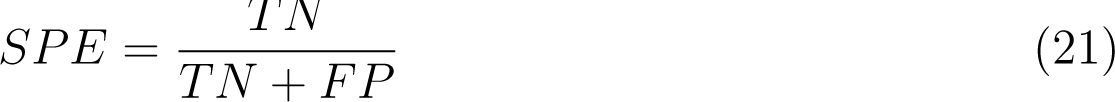

where the number of true positives, true negatives, false positives, and false negatives are denoted by TP, TN, FP, and FN, respectively. When calculating the SEN and SPE of a specific functional peptide, that class of peptide is considered a positive sample and other peptides without that function are considered negative samples.

### 4.4 Implementation details

Our proposed model was implemented by using PyTorch1.12 in a computing server equipped with Intel(R) Xeon(R) Gold 6248R CPU @ 3.00GHz and NVIDIA A100 GPU. The proposed models were trained in 100 epochs with batch size 32 and AdamW optimizer [22]. Since the pLM has already pre-trained, we set a small learning rate of 5e-5 to fine-tune it. While the learning rate of the GAT is 1e-3 and 5e-4 for the MFBP and MFTP datasets respectively. We constructed fully-connected graphs with 5 and 21 nodes for the MFBP and MFTP datasets respectively and both with the initialized edge weights of 1. The number of attention heads in GAT is 6 and the dimension of node features in each attention head is 128. The final node features in GAT have a dimension of 768, same to the representation obtained from the language model. The norm constraint *ɛ* of adversarial training is set to 0.5. In order to reduce the effects caused by the random initialisation of the deep learning framework and to maintain a consistent setting with the compared methods [33, 44], all model results were trained 10 times repeatedly and the prediction results were averaged as the final prediction for testing samples.

### 4.5 Antibacterial and anticancer experiment setup

#### Peptide synthesis

The peptides used in this study were synthesized by solid-phase peptide synthesis by Beijing Liuhe Bada Gene Technology Co., LTD, and their accurate molecular weights were determined by mass spectrometry. The purity of all peptides was determined by high-performance liquid chromatography, and all purity was greater than 90%.

#### Bacterial inhibition experiment

S. aureus bacteria strain was streaked on Luriae-Bertani (LB) agar medium and incubated at 37°C overnight. The individual colony was picked into LB culture medium and shaken at 120 r.p.m. at 37°C overnight. The LB bacterial suspension was diluted to the predetermined starting concentration (optical density at 600 nm (OD600) = 0.1) and then again diluted 1,000 times for the inhibition test. We thawed freeze-dried powder of peptides and dissolved in double-distilled water to 50 mmolL*^−^*^1^. We set three experimental groups to test peptide antibacterial activity: (1) blank control group, 50 *µ*L of LB solution; (2) bacterial control group, 25 *µ*L of LB solution and 25 *µ*L of bacterial solution; and (3) peptide group (500 *µ*molL*^−^*^1^), 23 *µ*L of LB solution, 50 *µ*L of bacterial solution and 2 *µ*L of peptide solution. Experiments were performed on 96-well plates with each single well containing 50 *µ*L of final volume. After culture at 37°C for 12 hours, the absorbance value of each well was determined by using a microplate reader at 600 nm. All experiments were performed with three independent replicates.

#### Antitumor inhibition experiment

Antitumor inhibition of peptides was determined using cytotoxicity assay for T24, Hela, HepG2 cell lines. Exponentially growing cells were seeded in 96-well microtiter plates at a density of approximately 5 10^3^ cells per well. After 24 hours incubation at 37°C with 5% CO_2_ in atmosphere, medium was replaced with fresh medium, and peptides at final concentrations of 500 *µ*gmL*^−^*^1^ each were added, followed by 24 hours incubation. Cell viability was monitored by the addition of 5 mgmL*^−^*^1^ 3-[4,5-dimethylthiazol-2-yl]-2,5-diphenyltetrazolium bromide (MTT) solution and measurement of the absorbance value at OD570 after incubated for 4 hours. Each peptide experiment was performed with three replicates. The optical density of wells containing cells cultured without peptides was assumed to represent 100% cell viability.

## Supplementary Files

### Figures

**Figure S1:**
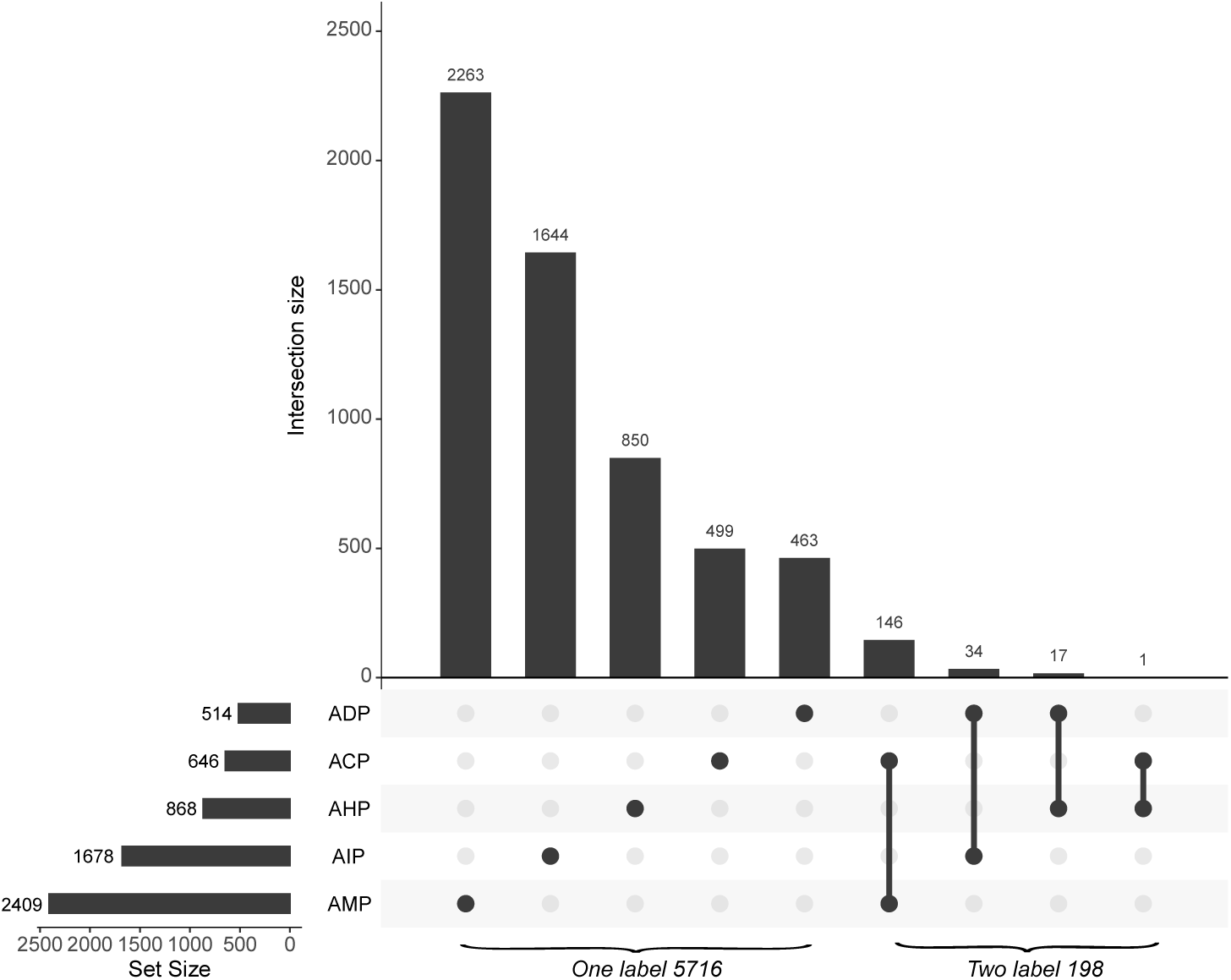
The distribution of multi-functional bioactive peptides in MFBP dataset. The vertical bar on the upper side shows the size of mono-functional and multi-functional peptide categories. The functions of each category are marked with the dots. The bottom horizontal bar shows the size of peptides in each function set.

**Figure S2:**
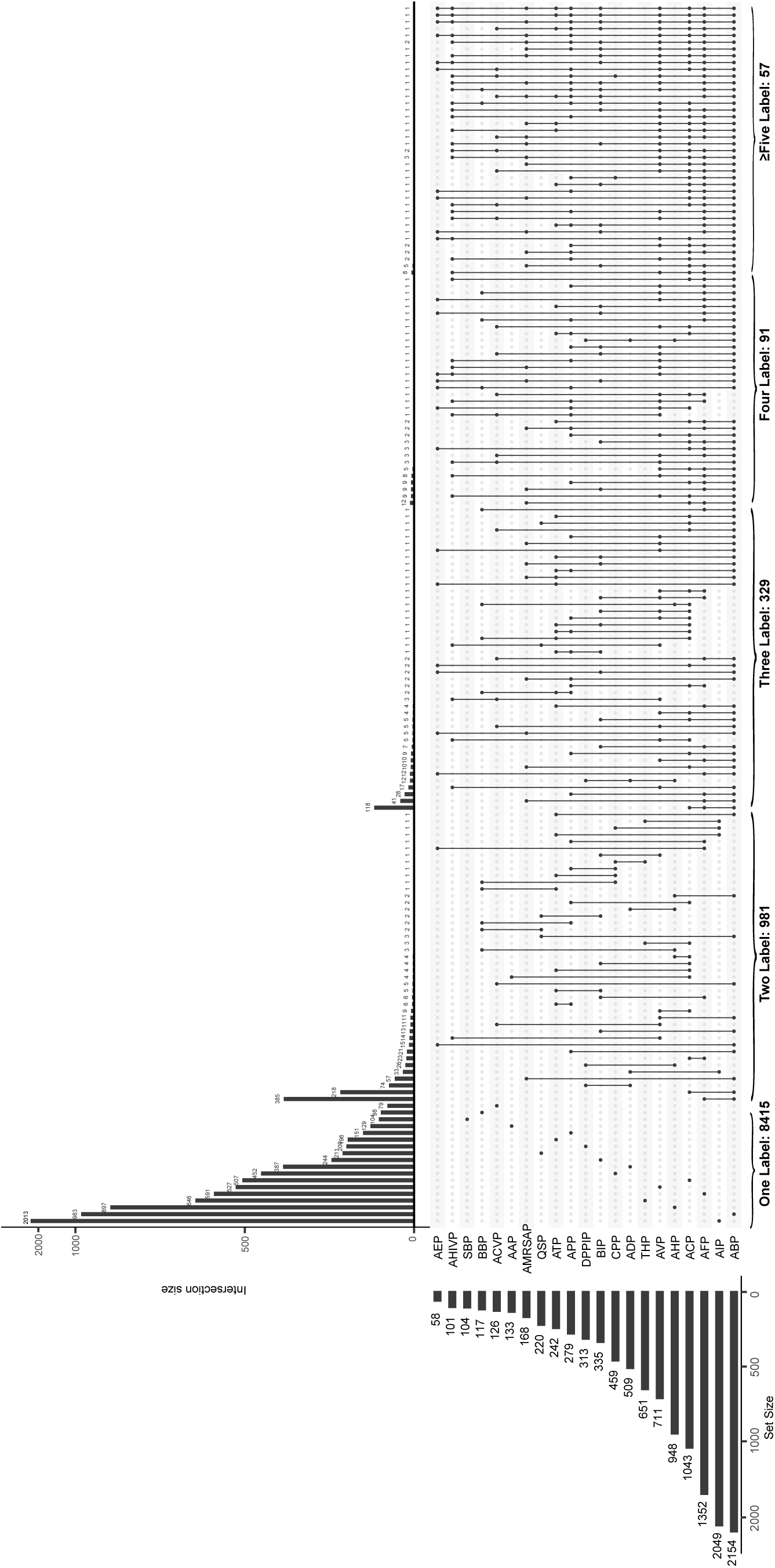
The distribution of multi-functional therapeutic peptides in MFTP dataset (zoom in to see the details). The horizontal bar on the left side shows the size of mono-functional and multi-functional peptide categories. The functions of each category are marked with the dots. The vertical bar shows the size of peptides in each function set.

**Figure S3:**
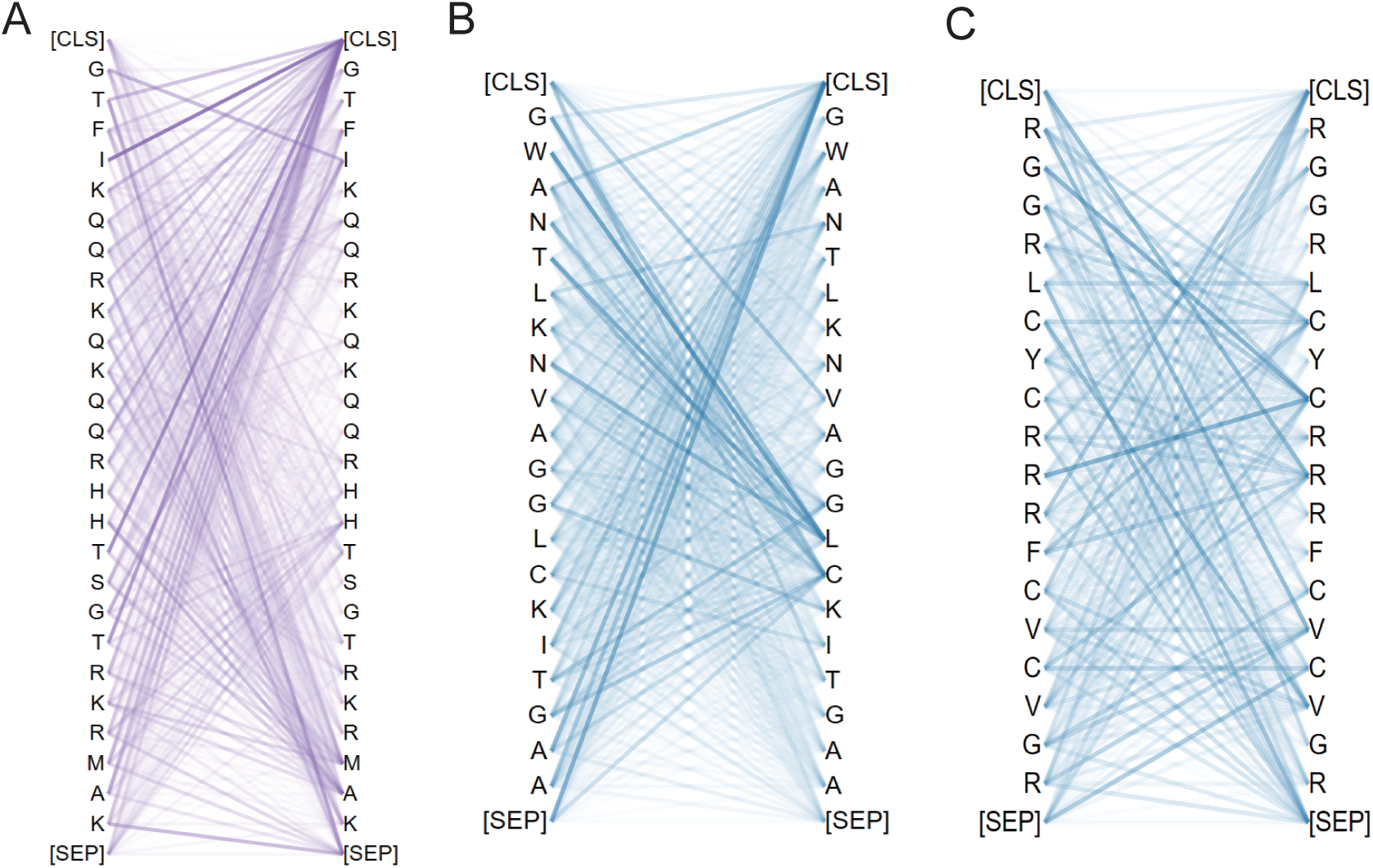
The 3 AMP cases to show attention weights between amino acids in a peptide sequences visualized via bertviz. The darker colors represent higher attention scores.

**Figure S4:**
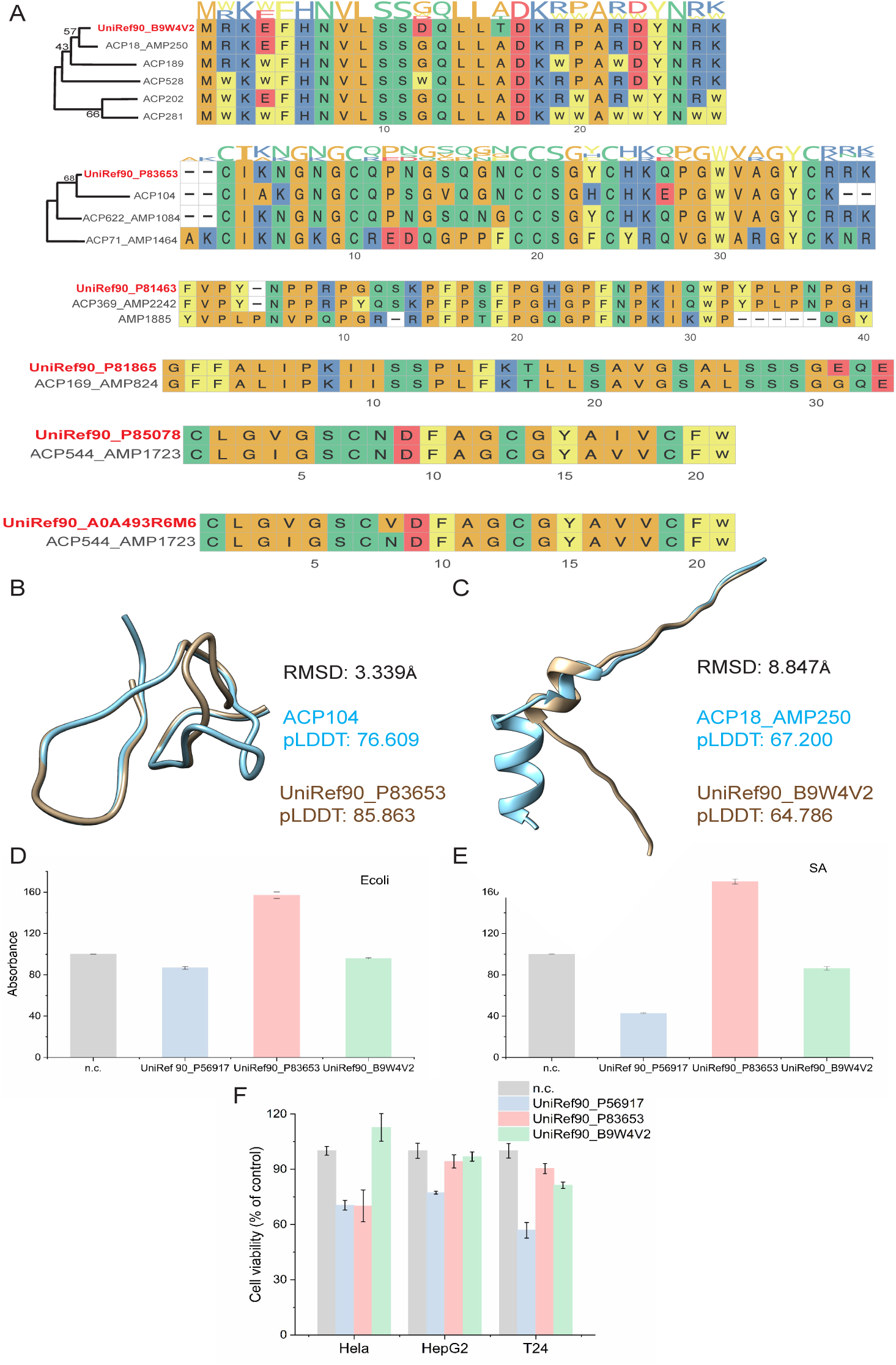
(A) MSA and pylogenetic trees of candidate peptides. (B-C) Structure alignment of homologous sequences with the closest evolutionary distance in pylogenetic tree. (D-E) Bacterial inhibition effect of peptides UniRef90 P56917, UniRef90 P83653 and UniRef90 B9W4V2 on E. coli and S. aureus bacteria strains. (F) Cytotoxic effect of peptides UniRef90 P56917, UniRef90 P83653 and UniRef90 B9W4V2 on T24, Hela, HepG2 cells. Peptides at 500 *µ*gmL*^−^*^1^ were added to cell culture for 24 hours, followed by MTT cytotoxicity assay. The results are the average of three independent replicates.

### Tables

**Table S1:** The performance comparison of different feature extraction methods with and without GAT on MFBP dataset. The best values are highlighted in bold.

| Method | Precision ↑ | Coverage ↑ | Accuracy ↑ | Absolute true ↑ | Absolute false ↓ |
| --- | --- | --- | --- | --- | --- |
| CF | 0.623 | 0.619 | 0.606 | 0.576 | 0.118 |
| with GAT | 0.667 | 0.649 | 0.646 | 0.623 | 0.121 |
| CNN | 0.688 | 0.706 | 0.687 | 0.667 | 0.108 |
| with GAT | 0.726 | 0.745 | 0.726 | 0.706 | 0.102 |
| RNN | 0.741 | 0.747 | 0.732 | 0.709 | 0.094 |
| with GAT | 0.759 | 0.763 | 0.747 | 0.719 | 0.094 |
| CNN-BiLSTM | 0.735 | 0.749 | 0.734 | 0.718 | 0.095 |
| with GAT | 0.761 | 0.775 | 0.757 | 0.735 | 0.091 |
| pLM | 0.774 | 0.784 | 0.774 | 0.762 | 0.082 |
| **with GAT** | **0.777** | **0.785** | **0.776** | **0.767** | **0.082** |

Note: ↑ means a larger value is better on this metric; ↓ means a smaller value is better on this metric.

**Table S2:** The performance comparison of different feature extraction methods with and without GAT on MFTP dataset. The best values are highlighted in bold

| Method | Precision ↑ | Coverage ↑ | Accuracy ↑ | Absolute true ↑ | Absolute false ↓ |
| --- | --- | --- | --- | --- | --- |
| CF | 0.362 | 0.333 | 0.323 | 0.283 | 0.047 |
| with GAT | 0.421 | 0.408 | 0.381 | 0.324 | 0.049 |
| CNN | 0.615 | 0.603 | 0.572 | 0.509 | 0.037 |
| with GAT | 0.637 | 0.626 | 0.592 | 0.525 | 0.037 |
| RNN | 0.624 | 0.614 | 0.584 | 0.522 | 0.035 |
| with GAT | 0.653 | 0.645 | 0.610 | 0.542 | 0.035 |
| CNN-BiLSTM | 0.661 | 0.650 | 0.616 | 0.546 | 0.033 |
| with GAT | 0.680 | 0.671 | 0.635 | 0.564 | 0.034 |
| pLM | 0.709 | 0.705 | 0.667 | 0.595 | 0.032 |
| **with GAT** | **0.721** | **0.722** | **0.697** | **0.605** | **0.032** |

Note: ↑ means a larger value is better on this metric; ↓ means a smaller value is better on this metric.

**Table S3:** The performance comparison of our proposed method iMFP-LG with the state-of-the-art methods on the MFBP dataset. The best values are highlighted in bold.

| Method | Precision ↑ | Coverage ↑ | Accuracy ↑ | Absolute true ↑ | Absolute false ↓ |
| --- | --- | --- | --- | --- | --- |
| CLR [33] | 0.667 | 0.677 | 0.666 | 0.655 | 0.133 |
| RAKEL [33] | 0.649 | 0.648 | 0.648 | 0.647 | 0.141 |
| MLDF [33] | 0.649 | 0.649 | 0.648 | 0.646 | 0.119 |
| RBRL [33] | 0.650 | 0.651 | 0.649 | 0.646 | 0.140 |
| MLBP [33] | 0.710 | 0.720 | 0.709 | 0.697 | 0.106 |
| MPMABP [19] | 0.728 | 0.749 | 0.727 | 0.704 | 0.101 |
| **iMFP-LG** | **0.797** | **0.803** | **0.796** | **0.788** | **0.078** |

Note: ↑ means a larger value is better on this metric; ↓ means a smaller value is better on this metric.

**Table S4:** The performance comparison of our proposed method iMFP-LG with the state-of-the-art methods on the MFTP dataset. The best values are highlighted in bold.

| Method | Precision ↑ | Coverage ↑ | Accuracy ↑ | Absolute true ↑ | Absolute false ↓ |
| --- | --- | --- | --- | --- | --- |
| BR [44] | 0.427 | 0.437 | 0.394 | 0.325 | 0.050 |
| CLR [44] | 0.418 | 0.428 | 0.387 | 0.320 | 0.047 |
| RAKEL [44] | 0.349 | 0.317 | 0.307 | 0.265 | 0.052 |
| RBRL [44] | 0.513 | 0.507 | 0.478 | 0.426 | 0.057 |
| PrMFTP [44] | 0.699 | 0.669 | 0.651 | 0.593 | **0.031** |
| **iMFP-LG** | **0.730** | **0.730** | **0.689** | **0.616** | 0.032 |

Note: ↑ means a larger value is better on this metric; ↓ means a smaller value is better on this metric.

**Table S5:** The performance of the MFBP experiment model with 10 repetitions on MFBP test set.

| Model | Precision ↑ | Coverage ↑ | Accuracy ↑ | Absolute true ↑ | Absolute false ↓ |
| --- | --- | --- | --- | --- | --- |
| Model0 | 0.778 | 0.792 | 0.777 | 0.761 | 0.090 |
| Model1 | 0.789 | 0.805 | 0.788 | 0.769 | 0.085 |
| Model2 | 0.774 | 0.784 | 0.773 | 0.760 | 0.089 |
| Model3 | 0.770 | 0.776 | 0.769 | 0.762 | 0.091 |
| Model4 | 0.776 | 0.787 | 0.775 | 0.761 | 0.090 |
| Model5 | 0.758 | 0.767 | 0.757 | 0.747 | 0.094 |
| Model6 | 0.788 | 0.801 | 0.787 | 0.772 | 0.085 |
| Model7 | 0.776 | 0.786 | 0.774 | 0.761 | 0.090 |
| Model8 | 0.773 | 0.783 | 0.771 | 0.759 | 0.091 |
| Model9 | 0.761 | 0.768 | 0.760 | 0.751 | 0.092 |
| Model<sub>avg</sub> | 0.797 | 0.803 | 0.796 | 0.788 | 0.078 |

Note: Model0-9 means the results of the model repeated 10 times with different random seeds. Model<sub>avg</sub> means averaged the results of 10 model (Model0-9) predictions as the final prediction for testing samples.

**Table S6:** The performance of the MFTP experiment model with 10 repetitions on MFTP test set.

| Model | Precision ↑ | Coverage ↑ | Accuracy ↑ | Absolute true ↑ | Absolute false ↓ |
| --- | --- | --- | --- | --- | --- |
| Model0 | 0.695 | 0.713 | 0.657 | 0.575 | 0.039 |
| Model1 | 0.695 | 0.712 | 0.657 | 0.575 | 0.038 |
| Model2 | 0.693 | 0.713 | 0.660 | 0.582 | 0.038 |
| Model3 | 0.688 | 0.701 | 0.652 | 0.575 | 0.037 |
| Model4 | 0.698 | 0.716 | 0.660 | 0.575 | 0.039 |
| Model5 | 0.695 | 0.713 | 0.658 | 0.574 | 0.039 |
| Model6 | 0.711 | 0.728 | 0.674 | 0.593 | 0.037 |
| Model7 | 0.707 | 0.713 | 0.662 | 0.582 | 0.037 |
| Model8 | 0.702 | 0.717 | 0.665 | 0.588 | 0.037 |
| Model9 | 0.700 | 0.705 | 0.658 | 0.581 | 0.038 |
| Model<sub>avg</sub> | 0.730 | 0.730 | 0.689 | 0.616 | 0.032 |

Note: Model0-9 means the results of the model repeated 10 times with different random seeds. Model<sub>avg</sub> means averaged the results of 10 model(Model0-9) predictions as the final prediction for testing samples.

**Table S7:** The settings of STREME for finding the motif of AMP in MFBP dataset.

| Parameter | value |
| --- | --- |
| Minimum width | 3 |
| Maximum width | 6 |
| $p$ value threshold | 0.005 |
| Number of motifs | 10 |
| Align sequences on their | Centers |

